# Developmental exposure to polychlorinated biphenyls prevents recovery from noise-induced hearing loss and disrupts the functional organization of the inferior colliculus

**DOI:** 10.1101/2023.03.23.534008

**Authors:** Baher A. Ibrahim, Jeremy Louie, Yoshitaka Shinagawa, Gang Xiao, Alexander R. Asilador, Helen J. K. Sable, Susan L. Schantz, Daniel A. Llano

**Affiliations:** Department of Molecular & Integrative Physiology, the University of Illinois Urbana-Champaign, Urbana, IL 61801, USA; Beckman Institute for Advanced Science & Technology, the University of Illinois Urbana-Champaign, Urbana, IL 61801, USA; Neuroscience Program, the University of Illinois Urbana-Champaign, Urbana, IL 61801, USA; Department of Comparative Biosciences, the University of Illinois Urbana-Champaign, Urbana, IL 61801, USA; Carle Illinois College of Medicine, the University of Illinois Urbana-Champaign, Urbana, IL 61801, USA; The Department of Psychology, The University of Memphis, Memphis, TN 38152, USA

## Abstract

Exposure to combinations of environmental toxins is growing in prevalence, and therefore understanding their interactions is of increasing societal importance. Here, we examined the mechanisms by which two environmental toxins – polychlorinated biphenyls (PCBs) and high-amplitude acoustic noise – interact to produce dysfunction in central auditory processing. PCBs are well-established to impose negative developmental impacts on hearing. However, it is not known if developmental exposure to this ototoxin alters the sensitivity to other ototoxic exposures later in life. Here, male mice were exposed to PCBs in utero, and later as adults were exposed to 45 minutes of high-intensity noise. We then examined the impacts of the two exposures on hearing and the organization of the auditory midbrain using two-photon imaging and analysis of the expression of mediators of oxidative stress. We observed that developmental exposure to PCBs blocked hearing recovery from acoustic trauma. In vivo two-photon imaging of the inferior colliculus revealed that this lack of recovery was associated with disruption of the tonotopic organization and reduction of inhibition in the auditory midbrain. In addition, expression analysis in the inferior colliculus revealed that reduced GABAergic inhibition was more prominent in animals with a lower capacity to mitigate oxidative stress. These data suggest that combined PCBs and noise exposure act nonlinearly to damage hearing and that this damage is associated with synaptic reorganization, and reduced capacity to limit oxidative stress. In addition, this work provides a new paradigm by which to understand nonlinear interactions between combinations of environmental toxins.

**Significance statement:** Exposure to common environmental toxins is a large and growing problem in the population. This work provides a new mechanistic understanding of how the pre-and postnatal developmental changes induced by polychlorinated biphenyls could negatively impact the resilience of the brain to noise-induced hearing loss later in adulthood. The use of state-of-the-art tools, including in vivo multiphoton microscopy of the midbrain helped in identifying the long-term central changes in the auditory system after the peripheral hearing damage induced by such environmental toxins. In addition, the novel combination of methods employed in this study will lead to additional advances in our understanding of mechanisms of central hearing loss in other contexts.

## Introduction

Overexposure to occupational noise is considered one of the main factors negatively impacting hearing in the USA and Europe and is growing in prevalence (Bergström and Nyström, 1986; Daniell et al., 2006; Ding et al., 2019). Contamination of the environment with ototoxic chemicals could also exacerbate the effect of noise-induced hearing loss (NIHL). Polychlorinated biphenyls (PCBs) are associated with hearing loss and other health deficits (Powers et al., 2006; Powers et al., 2009; Min et al., 2014) (Wu et al., 1984; Safe, 1993; Goldey et al., 1995; Morse et al., 1996; Brouwer et al., 1999; Xie et al., 2019). PCBs were banned by the U.S. Environment Protection Agency (Ross, 2004) but are highly chemically stable, which has led to widespread and persistent environmental contamination (Beyer and Biziuk, 2009). Given that PCBs can enter the placenta and breast milk (Jacobson et al., 1984), they also pose a developmental threat (Guo et al., 2004) including to the cochlea (Uziel, 1986; Goldey et al., 1995; Wong et al., 1997; Knipper et al., 2000; Song et al., 2008) and developmental exposure to PCBs impairs the hearing of humans and animals (Goldey et al., 1995; Herr et al., 1996; Crofton and Rice, 1999; Crofton et al., 2000; Lasky et al., 2002; Powers et al., 2006; Kenet et al., 2007; Powers et al., 2009; Jusko et al., 2014; Min et al., 2014; Palkovičová Murínová et al., 2016; Sadowski et al., 2016; Lee et al., 2021). Given the high population prevalence of dual exposure to PCBs and noise, the objective of the current study was to examine the interaction between developmental exposure to PCBs and NIHL later in adulthood in both the peripheral and central auditory systems. We, therefore, exposed mice to PCBs and noise overexposure similar to the sequence that also occurs in humans: prenatal + breastmilk exposure to PCBs followed by adult overexposure to noise. Given previous work showing changes in inhibitory tone in the major auditory integration center in the midbrain, the inferior colliculus (IC), after either PCB or noise overexposure (Abbott et al., 1999; Dong et al., 2010b; Dong et al., 2010a; Auerbach et al., 2014; Poon et al., 2015; Bandara et al., 2016; Sadowski et al., 2016; Knipper et al., 2021; Lee et al., 2021), we examined inhibitory neuronal activity in the IC of adult mice exposed to PCBs perinatally, noise in adulthood, or both, using two-photon imaging of the IC *in vivo*. In addition, given previous work showing that pathological disruptions of inhibition may be related to oxidative stress (as reviewed (Ibrahim and Llano, 2019)), we measured markers of oxidative stress in the IC. We observed that the developmental exposure to PCBs impaired the hearing of the male mice to low frequency pure tones. While developmental exposure to PCBs did not exacerbate the NIHL, it blocked the hearing recovery one week from acoustic trauma at the same low frequencies. Using two-photon imaging, developmental exposure to PCBs and high level of noise later in adulthood showed a disruption of the tonotopic maps of the dorsal cortex of the IC (DCIC), which was characterized with a wide non-responsive zone to acoustic stimulation, increase in the stimulus level required to evoke the cellular activity of the responsive cells, and the downregulation of inhibition, which was associated with low resilience to oxidative stress. These results suggest nonlinear interactions between the developmental exposure to PCBs and NIHL at the level of both peripheral and central auditory system.

## Materials and methods

### Animals

Wild-type female Swiss Webster (SW) mice (Jackson lab, # 000689), approximately three months of age, were used for the initial PCBs dosing. The female mice were bred with male GAD67-GFP knock-in mice of the same SW background (developed and shared with permission from Dr. Yuchio Yanagawa at Gunma University and obtained from Dr. Douglas Oliver at the University of Connecticut) to produce offspring where GFP is exclusively expressed in the GABAergic cells (Tamamaki et al., 2003) to visualize and distinguish the GABAergic cells from non-GABAergic cells in the offspring. All procedures were performed following protocol #19104, which was approved by the Institutional Animal Care and Use Committee (IACUC) at The University of Illinois at Urbana-Champaign. The mice were individually housed in standard plastic cages with corncob bedding, in a temperature- and a humidity-controlled room that was maintained on a 12-h light/dark cycle (lights off at 1900 h). Food and tap water were available ad libitum.

### PCB exposure

The timeline of the experimental procedures is shown in figure 1A. Given that eating PCB-contaminated fish was reported to be one of the main routes of human exposure to PCBs (Fitzgerald et al., 1996; Fitzgerald et al., 1998), the Fox River PCB mixture, which mimics the PCB levels in contaminated Walleye fish from the Fox River of Northeast Wisconsin, was used in this study as previously done on rats (Powers et al., 2006; Sadowski et al., 2016; Lee et al., 2021). This PCB mixture consisted of 35% Aroclor 1242, 35% Aroclor 1248, 15% Aroclor 1254, and 15% Aroclor 1260 (Kostyniak et al., 2005). In study phase one, the developmental toxicity of PCBs on hearing was examined in mice. The female mice were randomly assigned to three different groups receiving daily PCBs doses of 0, 6, or 12 mg/kg body weight. A dose of 6 mg/kg PCBs was reported to impair hearing in rats compared to lower doses (Powers et al., 2006). Exposure began 28 days before mating and continued until pups were weaned on postnatal day 21 (P21). The PCBs mixture was diluted in corn oil (Mazola®) and pipetted onto 40-50 mg of vanilla wafer cookie (Nilla Vanilla Wafers®) at a volume of 0.4 ml/kg. At the dosing time, each female mouse was taken to a separate empty plastic cage (with no bedding to easily identify the cookie pellet) at approximately 1400 h. The PCBs-contaminated cookie was placed in the center of its dosing cage allowing the animal to fully eat it. Aligned with the previous report, it was found that the average time for the mouse to fully eat the cookie was 30 minutes (Cadot et al., 2012). The same dosing paradigm was followed in study phase II, which examined the interaction between developmental exposure to PCBs and noise exposure in adulthood.

**Figure 1.**
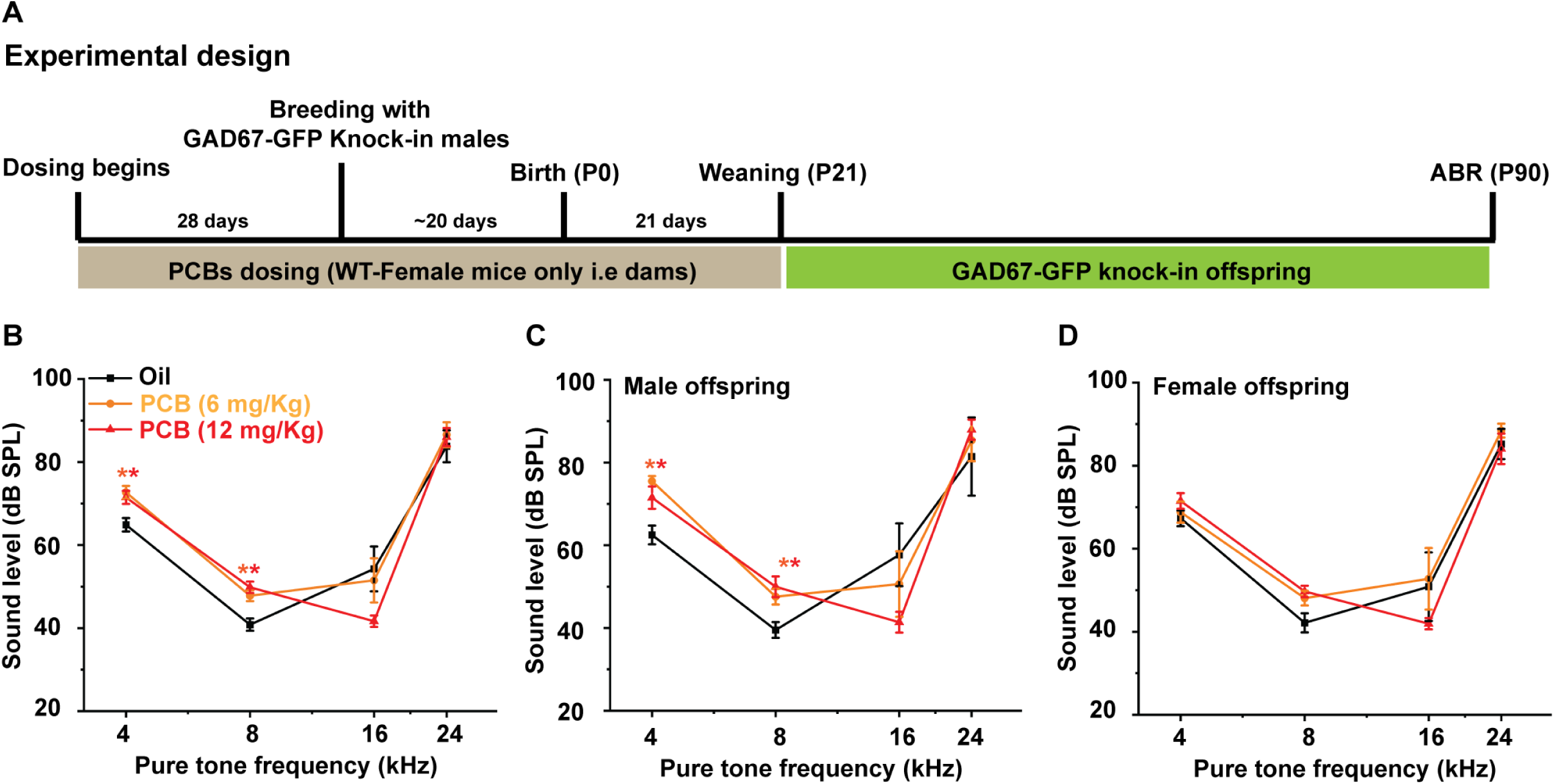
Developmental exposure to PCBs impairs hearing in male mice. ***A,*** The timeline of the experimental design of the PCB dosing of dams and hearing assessment of the pups. ***B,*** A line plot of the hearing threshold across different frequencies for all pups came from dams under different exposures (One-Way ANOVA: f (2,33) = 6.1, p = 0.005, Fisher post hoc test: *p = 0.002 and 0.007 for Oil vs. 6 and 12 mg/kg PCB, respectively, and p = 0.58 for 6 vs. 12 mg/kg PCB at 4 kHz & f (2,33) = 10.8, *p = 2.4×10^-4^ and 7.1×10-5 for Oil vs. 6 and 12 mg/kg PCB, respectively, and p = 0.28 for 6 vs. 12 mg/kg PCB at 8 kHz & f _(2,32)_ = 3.1, p = 0.06 at 16 kHz & f _(2,26)_ = 1.2, p = 0.31 at 24 kHz). ***C,*** A line plot of the hearing threshold across different frequencies for male pups came from dams under different exposures (One-Way ANOVA: f _(2,16)_ = 8.3, p = 0.003, Fisher post hoc test: *p = 9.1×10^-4^ and 0.01 for Oil vs. 6 and 12 mg/kg PCB, respectively, and p = 0.19 for 6 vs. 12 mg/kg PCB at 4 kHz & f _(2,16)_ = 5.5, p = 0.02, Fisher post hoc test: *p = 0.02 and 0.005 for Oil vs. 6 and 12 mg/kg PCB, respectively, and p = 0.43 for 6 vs. 12 mg/kg PCB at 8 kHz & f _(2,15)_ = 2.0, p = 0.17 at 16 kHz & f _(2,11)_ = 1.1, p = 0.36 at 24 kHz). **D,** A line plot of the hearing threshold across different frequencies or female pups came from dams under different exposures (One-Way ANOVA: f _(2,14)_ = 1.1, p = 0.36 at 4 kHz & f _(2,13)_ = 3.2, p = 0.08 at 8 kHz & f _(2,14)_ = 1.1, p = 0.35 at 16 kHz & f _(2,14)_ = 0.4, p = 0.67 at 24 kHz). *: The significance against Oil treated group.

### Breeding

In all study phases, each female mouse was paired with an unexposed GAD67-GFP male mouse of a SW background. After breeding, the females were monitored once daily for the presence of a sperm plug (i.e., gestational day 0), and mice were paired for 15 consecutive days regardless of whether a sperm plug was observed. The pups at day 7 of age were phenotyped for the presence of GFP under a fluorescence microscope via transcranial imaging (Tamamaki et al., 2003). This process was done by gently handling and placing each pup under a fluorescence microscope. GFP-positive pups were detected by examining the green fluorescent signals of the GFP (excitation: 472/30 nm, dichroic 505 nm, emission 520/35 nm long pass). Then, only GFP-positive pups were kept. After weaning, one male and one female from each litter were randomly selected to serve as subjects for this study. The pups were housed in same-exposure and same-sex cages of up to 5 mice/cage.

### Noise exposure

In study phase two, the paradigm of noise exposure was selected to induce a TTS (Amanipour et al., 2018). Whether the animals were developmentally exposed to PCBs or not, the three-month-old animals were all exposed to 110 dB SPL broadband noise for 45 minutes (Fig. 2A. The noise was delivered by Fostex super tweeter (T925A, Fostex Crop, Japan) using a pre-amplifier (SLA1, Studio linear amplifier). The noise exposure was done under anesthesia induced with a mixture of ketamine hydrochloride (100 mg/kg), xylazine (3 mg/kg), and acepromazine (3 mg/kg) to avoid the possible occurrence of audiogenic seizures as shown before in rats (Poon et al., 2015; Bandara et al., 2016). The exposure was done in a sound-attenuated dark room and was supervised from outside using an IR camera (ELP HD digital camera). The anesthesia was maintained during noise exposure using only ketamine (100 mg/kg).

**Figure 2.**
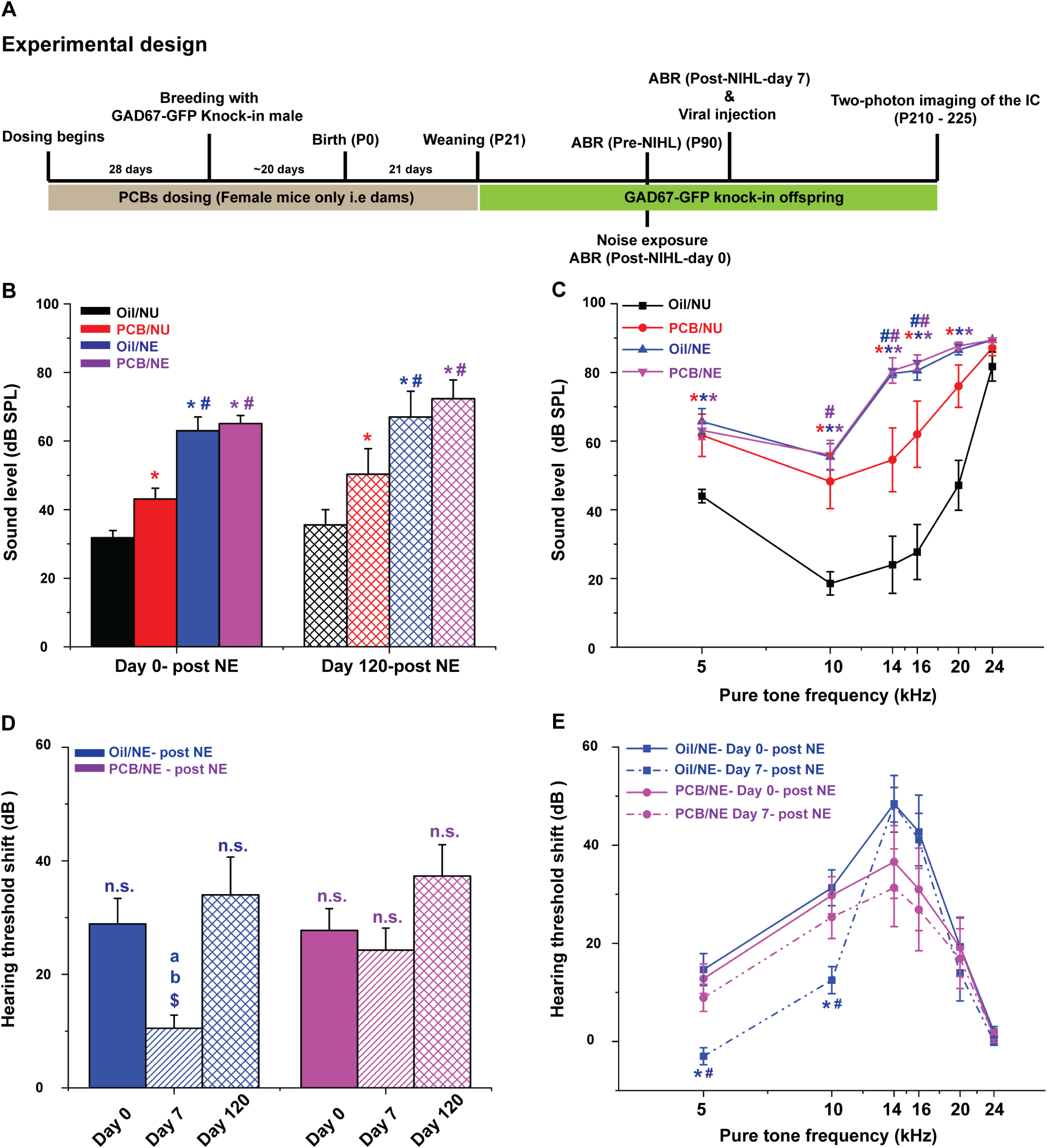
Developmental exposure to PCB inhibits the recovery from NIHL. ***A,*** The timeline of the experimental design of the PCBsdosing of dams, the overexposure to the noise of the pups, and hearing assessment of the pups before or after noise overexposure. ***B,*** A bar graph showing the threshold to flat noise under different exposures across two-time points (Day 0 and 120) (One-Way ANOVA: f _(7, 76)_ = 14.3, p = 9.6×10^-12^, Fisher post hoc test: *p = 0.009, 2.8×10^-7^, or 2.1×10^-8^ for Oil/NU vs. PCB/NU, Oil/NE, or PCB/NE, respectively, at day 0 & *p = 0.03, 9.3×10^-6^, or 3.9×10^-7^ for Oil/NU vs. PCB/NU, Oil/NE, or PCB/NE, respectively, at day 120 & ^#^p = 4.8×10^-4^ or 0.7.1×10^-5^ for PCB/NU vs. Oil/NE or PCB/NE, respectively, at day 0 & ^#^p = 0.03 or 0.005 for PCB/NU vs. Oil/NE or PCB/NE, respectively, at day 120 & p = 0.74 and 0.48 for Oil/NE vs. PCB/NE at day 0 and 120, respectively & p = 0.45, 0.24, 0.57, 0.30 for day 0 vs. 120 at Oil/NU, PCB/NU, Oil/NE, and PCB/NE, respectively). ***C***, A line graph of the ABR threshold to pure tones from different exposure groups (At 5 kHz, One-Way ANOVA: f _(3, 36)_ = 8.7, p = 1.8×10^-4^, Fisher post hoc test: *p = 0.004, 1.6×10^-4^, or 5.2×10^-5^ for Oil/NU vs.PCB/NU, Oil/NE, or PCB/NE, respectively & p = 0.19 for PCB/NU vs. Oil/NE, 0.11 for PCB/NU vs. PCB/NE, and 0.79 for Oil/NE vs. PCB/NE, (At 10 kHz, One-Way ANOVA: f _(3, 36)_ = 11.4, p = 2.0×10^-5^, Fisher post hoc test: *p = 0.005, 5.0×10^-5^, or 4.6×10^-6^ for Oil/NU vs.PCB/NU, Oil/NE, or PCB/NE, respectively & ^#^p = 0.02 PCB/NU vs.PCB/NE & p = 0.08 for PCB/NU vs. Oil/NE and 0.63 for Oil/NE vs PCB/NE), (At 14 kHz, One-Way ANOVA: f _(3, 36)_ = 15.1, p = 1.6×10^-6^, Fisher post hoc test: *p = 0.02, 4.0×10^-6^, or 1.2×10^-6^ for Oil/NU vs.PCB/NU, Oil/NE, or PCB/NE, respectively & ^#^p = 0.04 or 0.002 for PCB/NU vs.Oil/NE or PCB/NE, respectively & p = 0.81 for Oil/NE vs. PCB/NE), (At 16 kHz, One-Way ANOVA: f _(3, 34)_ = 12.6, p = 1.1×10^-5^, Fisher post hoc test: *p = 0.003, 1.7×10^-5^, or 3.5×10^-6^ for Oil/NU vs.PCB/NU, Oil/NE, or PCB/NE, respectively & ^#^p = 0.04 or 0.02 for PCB/NU vs.Oil/NE or PCB/NE, respectively & p = 0.81 for Oil/NE vs. PCB/NE), (At 20 kHz, One-Way ANOVA: f _(3, 36)_ = 11.8, p = 1.6×10^-5^, Fisher post hoc test: *p = 0.002, 1.4×10^-5^, or 7.6×10^-6^ for Oil/NU vs.PCB/NU, Oil/NE, or PCB/NE, respectively & p = 0.08 for PCB/NU vs. Oil/NE, 0.06 for PCB/NU vs. PCB/NE, and 0.94 for Oil/NE vs. PCB/NE), and (At 24 kHz, Kruskal-Wallis ANOVA, χ^2^= 9.04, p = 0.03, post hoc Dunn’s Test: p = 1.0, 0.22, and 0.41 for Oil/NU vs. PCB/NU, Oil/NE, and PCB/NE, respectively, and p = 1.0 for all remaining comparisons). ***D,*** A bar graph of the hearing threshold shift to the flat noise immediately, a week, or 4 months after acoustic trauma for the pups from dams exposed to PCBs or Oil (PCB/NE vs.Oil/NE) (One-Way ANOVA: f _(5, 38)_ = 4.4, p = 0.002, Fisher post hoc test: ^a^p = 0.003, p = 0.42, and ^b^p = 8.3×10^-4^ for day 0 vs. 7, day 0 vs. 120, and day 7 vs. 120, respectively, at Oil/NE & p = 0.58, 0.15, and 0.06 for day 0 vs. 7, day 0 vs. 120, and day 7 vs. 120, respectively, at PCB/NE & p = 0.85, ^$^p = 0.03, and p = 0.63 for Oil/NE vs PCB/NE at day 0, 7, and 120, respectively). ***E,*** A line graph of the hearing threshold shift to different pure tone frequencies immediately or a week after acoustic trauma for the pups from dams exposed to PCBs or Oil (PCB/NE vs.Oil/NE) (At 5 kHz, One-Way ANOVA: f _(3, 32)_ = 7.3, p = 7.0×10^-4^, Fisher post hoc test: *p = 1.5×10^-4^ and p = 0.32 for day 0 vs.7 at Oil/NE and PCB/NE, respectively & ^#^p = 0.007 for PCB/NE vs.Oil/NE at day 7), (At 10 kHz, One-Way ANOVA: f _(3, 32)_ = 4.9, p = 0.007, Fisher post hoc test: *p = 0.002 and p = 0.39 for day 0 vs.7 at Oil/NE and PCB/NE, respectively & ^#^p = 0.02 for PCB/NE vs.Oil/NE at day 7), and (One-Way ANOVA: f _(3, 32)_ = 1.7, p = 0.19 at 14 kHz, f _(3, 32)_ = 0.94, p = 0.42 at 16 kHz, f _(3, 32)_ = 0.17, p = 0.9 at 20 kHz, and f _(3, 32)_ = 0.51, p = 0.67 at 24 kHz). The exposure groups are plotted as Oil/NU: black, PCB/NU: red, Oil/NE: blue, and PCB/NE: purple; *: The significance against (Oil/NU) group; ^#^: The significance against (PCB/NU) group; ^$^: The significance against (Oil/NE) group, ^a^: The significance against day 0, and ^b^: The significance against day 120.

### ABR

As previously described (Lee et al., 2021), the ABR of three-month-old offspring was examined with or without noise exposure. In study phase one, the ABR was taken from the offspring at three months of age. In study phase two, the ABR was taken before (Pre-NIHL), immediately after (Post-NIHL-day 0), one week after (Post-NIHL-day 7), and four months after (Post-NIHL-day 120) noise exposure. ABRs were obtained at frequencies of 4, 5, 8, 10, 14, 16, 20, and 24 kHz as well as white noise. Animals were anesthetized with the same (ketamine hydrochloride, xylazine, and acepromazine) mixture, and the anesthesia was maintained during ABR measurements using only ketamine (100 mg/kg). Three subdermal electrodes were inserted [(one at the vertex (active electrode), one behind the right ear (reference electrode), and one behind the left ear (the ground electrode)]. The mouse was placed 5 cm from the speaker on a heating pad. Stimuli were presented using a Tucker-Davis (TDT) system 3 (Alachua, FL, USA), ES1 free-field speaker, with waveforms being generated by RPvdsEx software. The output of the TDT speaker was calibrated at all the relevant frequencies, using a PCB 377A06 microphone (Depew, NY, USA) and a Svantek 977 Sound & Vibration Analyser (Warsaw, Poland). Each frequency was presented for 5 ms (3 ms flat with 1 ms for both rise and fall times), at a rate of 21 Hz with a 45 ms analysis window. The ABR waveform was collected with TDT RA4PA Medusa Preamps with a 3 Hz −5 kHz filter and was averaged over 500 repeats for one sound stimulus. The visual inspection of the deflections of ABR within 10 ms from the stimulus’s onset continued until I-V waves were no longer present in the waveform. Although 86 dB SPL was the loudest level used, some animals were deaf at this level indicated by no identified ABR signals. To assign a numerical value for these deaf animals at 86 dB SPL sounds, a 90 dB SPL value was assumed as a threshold for those animals across different exposures. The reading of the ABR thresholds was conducted by an independent experimenter who was blind to the corresponding exposure groups of the animals. After ABR measurement, each animal was kept on a heating pad until a full recovery, then the animal was moved back to its home cage. After noise exposure, the hearing threshold shift was calculated on days 0, 7, and 120 indicated by [(Post-NIHL-day x) – (Pre-NIHL)], where x is the day after noise exposure i.e (0, 7, or 120).

### Viral injection

In study phase two, all mice across all groups were injected with AAV1.Syn.NES-jRGECO1a.WPRE.SV40 (100854-AAV1, Addgene) into the IC after the last ABR session (Post-NIHL-day 7). The virus drives the expression of the jRGECO1a in non-GABAergic and GABAergic cells via synapsin (*Syn*) promoter. The detailed surgical procedures were described before (Vaithiyalingam Chandra Sekaran et al., 2021). In brief, the surgery was done under aseptic techniques. Mice were firstly anesthetized intraperitoneally with the (ketamine hydrochloride, xylazine, and acepromazine) mixture. The anesthesia was maintained during the surgery using only ketamine (100 mg/kg). Then the mouse was placed into a Kopf Model 940 Small Animal Stereotaxic Instrument with a digital readout. After shaving and disinfecting the head with Povidone–iodide and 70% ethanol, a longitudinal incision was made in the scalp. The underneath surrounding tissue was injected with lidocaine (2% Covetrus, United States) intradermally as a local anesthetic. Also, the mice were injected with carprofen (3 mg/kg, Henry Schein Melville, NY, United States) subcutaneously as an analgesic for postoperative pain management. To protect the eyes from drying an ophthalmic ointment was applied. A small craniotomy was made over the IC targeting the DCIC using a surgical drill following these coordinates from lambda (x = 0.65 – 0.9 mm, y = 0.9 mm, and z = 0.4 mm). Glass micropipettes (3.5-inches, World Precision Instruments, Sarasota, FL) were pulled using a micropipette puller (P-97, Sutter Instruments, Novato, CA) and broken back to a tip diameter between 35 and 50 μm. The micropipette was filled with mineral oil (Thermo Fisher Scientific Inc, Waltham, MA) and attached to a pressure injector (Nanoliter 2010, World Precision Instruments, Sarasota, FL) connected to a pump controller (Micro4 Controller, World Precision Instruments, Sarasota, FL). The amount of the virus solution was then withdrawn by a pressure injector, and 450 nL of the viral solution with titer equal to 1.7 × 10^13^vgc/µL was injected into the DCIC at a flow rate (50 nl/minute). After the injection, the micropipette was kept in its place for another 5 minutes to avoid any solution retraction. The incision was then sutured using 5/0 thread-size, nylon sutures (CP Medical, Norcross, GA). After the surgery, the animals were kept on a heating pad for recovery. After awakening, animals were returned to their home cages.

### Craniotomy surgery for placing the cranial window and head post

The craniotomy was done four months after the viral injection, and the general procedures of craniotomy were previously described (Goldey et al., 2014), with modifications to place craniotomy over the IC. Before surgery, mice were anesthetized with the (ketamine hydrochloride, xylazine, and acepromazine) mixture. The anesthesia was maintained during the surgery and imaging using only ketamine (100 mg/kg). To prevent neural edema during or after the craniotomy, an intramuscular injection of dexamethasone sodium (4.8 mg/kg) was given just before the surgery using an insulin syringe. After placing the animal in the stereotaxic apparatus (David Kopf Instruments, USA), both eyes were protected by applying Opti care lubricant eye gel (Aventix Animal Health, Canada). The hair on the scalp was then removed by massaging the scalp with a depilatory cream (Nair) using a cotton-tipped applicator and leaving the cream on the scalp for 4-5 minutes. The cream was then removed by a thin plastic sheet (flexible ruler) to leave a hair-free area on the scalp. The remaining tiny hairs were then removed by alcohol swab and the area was then sterilized by applying 10% Povidone Iodine (Dynarex, USA) using a sterile cotton-tipped applicator. The medial incision was made with a scalpel blade #10, and 0.2 ml of 0.5% lidocaine was injected intradermally into the scalp. The skin extending from the medial line to the temporalis muscle was completely removed using a pair of micro scissors to produce a wide skinless area above the skull. A pair of no. 5/45 forceps were used to remove any remaining periosteum. The remaining dried or firmly attached pieces of periosteum were removed with a scalpel blade #10. The skull was cleaned with sterile saline and dried with gently pressurized air. Using the stereotaxic apparatus, a wide area of ∼3 x 4 mm above the left IC was made. A micro drill bit (Size #80, Grainger, USA) was used to drill through the skull starting from the rostro-lateral region to lambda. To prevent overheating of the superficial brain tissues and to mitigate the occasional spurts of skull bleeding during the drilling, ice-cold sterile saline was used to intermittently irrigate the surface. A stream of pressurized air was also applied during the drilling procedure to prevent overheating and remove the debris produced by the drilling. Caution was taken not to pierce the dura when performing the craniotomy while crossing the sagittal or the lambdoid sutures to avoid damaging the underlying sinuses. After drilling, the skull was irrigated in sterile saline and the bone flap (the undrilled bone over the craniotomy area) was gently examined for complete separation from the rest of the skull. Using a pair of no. 5/45 forceps, the bone flap was gently removed. To control the bleeding if it occurred, a piece of sterile hemostatic gel (Adsorbable Gelatin Sponge USP, Haemosponge, GBI, India) which was pre-soaked in ice-cold saline, was applied to the bleeding spot. Once the bleeding ceased, the brain was kept covered in sterile saline. In some surgeries, the dura was peeled off from the surface of the IC while removing the bone flap. In the surgeries where the dura remained intact, a Bonn microprobe (F.S.T, Germany, Item # 10032-13) was used to pierce the dura in an area that is not above the IC and devoid of cortical vasculature (e.g., a part of exposed cerebellum). After piercing, the dura was carefully and gently lifted, and a pair of no. 5/45 forceps were used to grab the dura to gently tear it to the extent of the transverse sinus to avoid bleeding. The cover glass was secured by a wooden trimmed piece of sterile cotton swab by gently pressing the cover glass from the top. A titanium head post as described before (Goldey et al., 2014) was glued carefully on the top of the skull to be at the same level as the cover glass following the manufacturer’s instructions, the C&B Metabond (Parkell, Japan).

### Two-photon imaging

Immediately after surgery, the anesthetized animal was taken and secured under the microscope objective by clamping the arms of the head post to two perpendicular metal posts mounted on the microscope stage. A custom-built 2P microscope was used. The optical and the controlling components were supplied from Bruker, Olympus, and Thorlabs. The imaging of the DC was made using a 20x water-immersion objective (LUMPlanFI/IR, 20X, NA: 0.95, WD: 2 mm; Olympus Corporation, Tokyo, Japan). Since the jRGECO1a calcium indicator was expressed in non-GABAergic and GABAergic neurons, and the GFP was only expressed in the GABAergic cells in the GAD67-GFP knock-in mouse, this was a good tool to distinguish the GABAergic (Green and red signals) from non-GABAergic cells (Red signals only). For imaging both the GFP or jRGECO1a signals, the excitation light was generated by InSight X3 laser (Spectra-Physics Lasers, Mountain View, CA, USA) tuned to a wavelength of 920 or 1040 nm, respectively. A layer of a 1:1 mixture of Multipurpose wavelengths ultrasound gel (National therapy, Canada) with double deionized water was used to immerse the objective. This gel was able to trap the water and reduce its evaporation during imaging. The emitted signals were detected by a photomultiplier tube (Hamamatsu H7422PA-4, Japan) following a t565lp dichroic and a Chroma barrier et525/70m filter for GFP and et595/50m filter for jRGECO1a signals. Images (512×512 pixels) were collected at a frame rate of 29.9 Hz at the resonant galvo mode. Data were collected from the dorsal surface of the IC by scanning the surface of the IC based on the GFP and jRGECO1a signals through the medial and lateral horizons of the IC. Generally, each field of view was selected based on the expression of jRGECO1a and being acoustically active using a search stimulus that was 500 ms broadband noise with zero modulation at 80 dB SPL. The frame timing of the scanner and the sound stimuli were both digitized and time-locked using a Digidata 1440A (Molecular Devices, Sunnyvale, CA, USA) with Clampex v. 10.3 (Molecular Devices, Sunnyvale, CA, USA).

### Acoustic stimulation

Using a custom-made MATLAB (The MathWorks, Natick, MA, USA) code, 500 ms pure tones were generated. Each pure tone is one of the thirty-five (5×7) combinations of sound pressure levels (80, 70, 60, 50,40 dB SPL) and carrier frequencies (5000-40000 Hz with a half-octave gap) that was presented with a cosine window. Each run is composed of these 35 combinations that were played in random sequence to the mice with a 600 ms interstimulus interval by a TDT RP2.1 processor (Tucker-Davis Technologies, US) and delivered by a TDT ES1 speaker (Tucker-Davis Technologies, US). Thus, stimuli were delivered every 1100 ms. Given the relatively fast kinetics of rGECO1a compared to earlier indicators (half decay time ∼300 ms (Dana et al., 2016), this interstimulus interval allowed for relatively dense sampling of stimulus space with minimal carryover of calcium waveforms between stimuli. Each animal was presented with different runs (mostly 9 runs on average). Each run has a different random sequence of the 35 combinations.

The output of the TDT ES1 speaker was calibrated using a PCB 377A06 microphone (Depew, NY, USA), which feeds SigCal tool to generate a calibration file for all tested frequencies (5-40 kHz). To enable the custom-made MATLAB code to read this calibration file, the values were first processed by MATLAB signal processing toolbox (sptool) to generate a 256-tap FIR filter to apply the calibration using the following parameters [arbitrary magnitudes, least square, order: 256, sampling rate: 97656.25, frequency vector (5-40 kHz), amplitude vector (40-80 dB SPL), and weight vector [ones (1,128)].

### Data processing

#### Data collection

The data were collected as separate movies (512×512 pixels) in a resonant galvo mode. Depending on the amplitude and frequency combinations for each type of acoustic stimulus, 40 seconds was assigned as a movie’s length for pure tone (35 stimulus combinations). Using ImageJ software (https://imagej.nih.gov/ij/), the z-projection was used to compute one single image representing either the sum, the standard deviation, or the median of all the image sequences in the movie. Based on these single images, the region of interest (ROI) was manually drawn around each detectable cell body.

#### Motion Correction and Filtering

The imread function from the OpenCV library was used in grayscale mode to import images into a numpy array from a single folder containing TIFF images. The array was then exported as a single TIFF stack into a temporary folder using the mimwrite function from the imageio library, and the process was repeated for each folder. The NoRMCorre algorithm (Pnevmatikakis and Giovannucci, 2017) embedded in the CaImAn library (Giovannucci et al., 2019) was used to apply motion correction to each of the TIFF files. The data were then imported into a numpy array, rounded, and converted to 16-bit integers. The images were filtered using a 2D Gaussian filter with a sigma value of 1 (Surface View) / 2 (Prism View) in each direction, then a 1D Gaussian temporal filter with a sigma value of 2 was applied using the ndimage.gaussian_filter and ndimage.gaussian_filter1d function from the scipy library, respectively.

#### Data Extraction

The ROI sets, which were manually created using ImageJ, were imported using the read_roi_zip function from the read_roi library. The sets were then used to create two masks; one mask was used as a replica of the ROIs and the second mask was made around the original ROI (roughly four times larger in the area). The smaller mask was applied to find the average pixel value within each ROI, while the larger mask was applied to find the average pixel value of the neuropil. The neuropil correction was applied using the following equation (Akerboom et al., 2012);

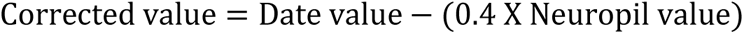

Δf/f values were then calculated by using the following equation.

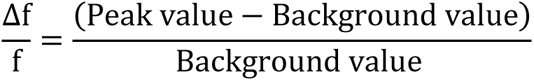

where the background value is the slope estimating the background levels with fluorescence bleaching factored in. The data were then reorganized so that all segments with the same stimulus frequency and stimulus amplitude were grouped.

#### Cell Flagging

The correlation coefficient between each of the trials was calculated using the stats. Pearsonr function from the scipy library. The average correlation coefficient was calculated for each stimulus frequency and stimulus amplitude. Similar to previous work (Wong and Borst, 2019), if the average correlation coefficient was above the threshold of 0.6, the cell was flagged as being responsive to that stimulus combination of frequency and amplitude, and the best frequency, which was defined as pure tone frequency that evoked the highest average response across all sound levels, was calculated for the cell (Barnstedt et al., 2015).

#### Tonotopic Map Generation

The average radius for all ROIs was calculated to ensure that all cells on the tonotopic map had uniform radii. A color key was also generated, with each shade of green representing one frequency. A blank canvas was generated using the Image module from the pillow library, and a circle was created for each cell and filled with the shade of green that represents the cell’s best frequency (Barnstedt et al., 2015). Cells that were non-responsive to stimuli were filled in red. The maps from all animals of the same group were aligned together based on the distance from the midline to construct a single map for each group. The tonotopic organization was determined by calculating the correlation between the distance (μm) of every cell from the most medial cell and its best frequency. The significant positive correlation is an indicator of the tonotopic organization when the cells are best tuned to lower and higher frequencies along the medial to the lateral axis. The statistics for the data set, such as the total number of cells, number of responsive cells, and number of non-responsive cells were tallied up and exported in an Excel spreadsheet.

#### Population Analysis

The average trace across all GABAergic cells and all non-GABAergic cells was plotted for each frequency and amplitude, similarly to the individual traces. The area under the curve (AUC) was used as a metric for calculation. For every animal, a heat map was generated within each cell type by normalizing all the values across the amplitude/frequency combinations to the highest value. Then one heat map was generated for each exposure group by averaging the normalized values at each amplitude/frequency combination. For every animal, the sound amplitude evokes 20% of the highest response was calculated as a threshold of the activity at each frequency. The (INH/EXC) ratio was calculated by dividing the AUC value of GABAergic cells over that of non-GABAergic cells at each amplitude/frequency combination for each animal across all exposure groups.

### Tissue preparation and Western Blot

#### Tissue preparation and protein quantification

Either after the last ABR measurement or immediately after the two-photon imaging, the animals, which were still under anesthesia, were decapitated and their brains were quickly isolated from the skull. The brains were then immediately immersed in isopentane solution, which was pre-cold using dry ice (Ibrahim and Briski, 2014; Alenazi et al., 2015). The frozen brains were then kept in the ultra-cold freezer (−80°C) until further processing. At the time of tissue extraction, the frozen brain was transferred to a cryostat (Leica) at −14° C and left for 5 minutes for equilibration. During this time, one mini tablet of protease and phosphatase Inhibitor (# A32959, Thermo Fisher) was added to 10 ml of ice-cold N-PER Neuronal Protein Extraction Reagent (# 87792, Thermo Fisher) by vigorous vortexing until the tablet was completely dissolved. The brain was then sectioned (50 μm) at the level of the IC. For each section that showed the IC, a micro puncher (EMS-Core Sampling, # 50-192-7735, Fisher Scientific) of 1 mm size, was used to dissect the tissue of the whole IC from the two hemispheres. All the dissected tissues from one animal were then dropped into the lysis solution (150 μl), which was then mechanically disrupted using an electrical homogenizer (Polytron, Switzerland). The homogenate was then centrifuged (10,000 x g for 10 minutes at 4°C), and the supernatant was then collected leaving the pellets behind. The protein concentration of the supernatant was then determined using Pierce™ BCA Protein Assay Kit (# PI23225, Fisher Scientific) according to the manufacturer’s instructions. The supernatant was then kept in the ultra-cold freezer (−80°C) until further processing.

#### Western Blotting

The protein of the lysates was then adjusted to 20 mg total protein using 4x Laemmli Sample Buffer (#1610747, BIO-RAD). The protein sample was denatured and run on 4-15% Mini-PROTEAN TGX Precast Protein Gels as per the manufacturer’s instructions (BIO-RAD). Blots were probed overnight at 4°C with antibodies recognizing GAD67 (1:1000, GAD1 (D1F2M) Rabbit mAb, #41318, Cell Signaling), SOD2 (1:1000, SOD2 (D9V9C) Rabbit mAb, #13194, Cell Signaling) and actin (1:1000, β-Actin Antibody #4967, Cell Signaling). Blots were then processed using anti-rabbit IgG, HRP-linked Antibody (#7074, Cell Signaling) followed by Super Signal West Pico PLUS Chemiluminescent Substrate (#34580, Thermo Fisher) as published before (Ibrahim et al., 2013).

### Statistics

Statistical analysis was done by Origin Pro 2022 software. The litter was the unit of the analysis, and the sex was nested within the litter. One litter from the control group was excluded from the study as the dam of this litter abandoned its pups resulting in the pups’ death. Also, not all litters have male and female pups, so the number of litters used in the analysis was different across the exposure groups and sexes. The normality of the data points within each group across the different sexes was examined by the Kolmogorov-Smirnov test. Based on the outcome, the parametric one-way ANOVA and Fisher post hoc tests were used for normal data, while the non-parametric Kruskal-Wallis ANOVA and Donn post hoc tests were used for the non-normal data to examine the significant differences between the exposure groups. A Chi-square test was used to examine the difference between the number of responsive vs. non-responsive cells to sound between each pair of groups. The significant effects were analyzed by the Fisher post hoc test, and the significance was called at p < 0.05 for all the statistical tests.

## Results

### Developmental exposure to PCBs impairs hearing in male mice

The toxic effect of developmental exposure to PCBs on hearing has been extensively studied in rats (Goldey et al., 1995; Herr et al., 1996; Crofton and Rice, 1999; Crofton et al., 2000; Lasky et al., 2002; Powers et al., 2006; Kenet et al., 2007; Powers et al., 2009; Sadowski et al., 2016; Lee et al., 2021), which do not provide a rich library of genetically modified models compared to mice. Examination of the toxic interaction between PCBs and noise on different cell types requires a genetically modified model that can aid in cell-type visualization. Therefore, mice were used in this study. To validate the mouse model for PCB studies, we first examined the toxic effect of PCBs on hearing in a mouse model that permits the examination of excitatory and inhibitory cell types. Given that fish are a primary source of PCBs exposure in humans (Persky et al., 2001; Judd et al., 2004; Roveda et al., 2006; Weintraub and Birnbaum, 2008), the Fox River PCB mixture (Sullivan et al., 1983) was used to simulate the human environmental exposure to PCBs. Wild-type female SW mice were daily given a 40-50 mg Nella cookie contaminated with either 0, 6, or 12 mg/kg of Fox River PCBs mixture (corn oil based) for 28 days before the breeding with PCBs-unexposed GAD67-GFP knock-in male mice (Tamamaki et al., 2003). Dosing continued throughout pregnancy and lactation until the weaning of their pups. ABR measurements were then taken from the three-month-old offspring to measure their hearing across different PCB developmental exposure levels. Under this experimental paradigm (Fig. 1A), the pre-and postnatal exposure to 6 or 12 mg/kg PCBs produced low-frequency (4 and 8 kHz) hearing impairment compared to vehicle-exposed dams (Fig. 1B), consistent with previous work (Powers et al., 2009). This hearing impairment was only significant in males (Figs. 1C and D). Therefore, only male offspring were selected for further experiments. There was no difference in the hearing threshold of the pups that came from dams either exposed to 6 or 12 mg/kg of PCBs (Fig. 1), which indicated the toxic effect of the PCBs on the hearing of the mice was not dose-dependent within the tested dose range. Therefore, the lower dose (6 mg/kg PCBs mixture) was used for the subsequent experiments.

### Developmental exposure to PCB inhibits the recovery from NIHL

After showing the developmental toxicity of PCB exposure on the hearing of male mice, we next examined the toxic interaction between PCB exposure during development and NIHL in adulthood. An additional cohort of wild-type SW female mice were dosed with 6 mg/kg Fox River PCB mixture daily for 28 days before breeding with PCBs-unexposed GAD67-GFP knock-in male mice and throughout pregnancy and lactation until the weaning of their pups (Fig. 2A). At three months of age, the GFP-positive male pups were randomly divided into two groups: a group exposed to 110 dB SPL broadband noise for 45 minutes (noise-exposed, NE) and a sham group (noise-unexposed, NU). Hence, a total of four subgroups, (Oil/NU, which is the control group of no PCBs or noise exposure), PCB/NU, Oil/NE, and PCB/NE. To measure the baseline hearing, an initial ABR was obtained from all offspring in all groups just before the noise or sham exposure at three months of age. To examine the acute toxic effect of noise exposure, another ABR was taken from all animals immediately after noise exposure (post-noise-day 0). This noise exposure paradigm was selected to induce a temporary threshold shift (Amanipour et al., 2018), so animals would regain their normal hearing one week after the acoustic trauma. Therefore, another ABR was taken from all animals one week after noise exposure (post-noise-day 7) to examine the hearing recovery of the animals after different exposures. Four months later, a final ABR was taken from all animals to assess any aging effect on the hearing baseline of the control group (Oil/NU) as well as the long-term effect of the other exposures on the peripheral auditory system. We observed that all adults that had been exposed to PCB and/or noise showed higher hearing thresholds than those of the control group (Oil/NU) to flat noise (Fig. 2B) and all pure tone frequencies below 24 kHz (Fig. 2C). However, the hearing impairment induced by noise exposure was significantly greater than that induced by PCBs indicated by higher hearing thresholds in Oil/NE or PCB/NE groups than PCB/NU to flat noise and the central frequencies (between 5 and 20 kHz) (Fig. 2B and C). Four months after the first ABR measurements, the hearing thresholds to flat noise of all groups did not change from those at day 0, which suggests any aging-related hearing effects across all groups are limited (Fig. 2B). Also, the difference between the thresholds of all groups was similar to the one measured at day 0 (Fig. 2B), which indicates that PCBs and/or noise exposures resulted in long-term permanent damage to the peripheral auditory system. Given the non-significant difference in the hearing threshold between the pups of the (Oil/NE) and (PCB/NE) groups for all tested stimuli (Fig. 2B and C), these data suggest that developmental exposure to a 6 mg/kg Fox River PCBs mixture did not exacerbate the acute effect of NIHL. The hearing recovery under different exposures was assessed by comparing the hearing threshold shift at different time points after noise exposure (Day 0, vs.7, and 120, see Methods). Initially, at day 0 or immediately after the noise exposure, the threshold shift of the animals of both Oil/NE and PCB/NE groups was increased indicating the acute damage of noise exposure to hearing. At day 7 or one week after noise exposure, while the threshold shift of the animals from the Oil/NE group was significantly reduced compared to that at day 0, the threshold shift of the animals from the PCB/NE group remained high indicating that the hearing recovery could be blocked by PCBs. At day 120 or four months after noise exposure, the threshold shift of the animals from the Oil/NE group recovered to be similar to that at day 0 or that of the animals from the PCB/NE group (Fig. 2D). The finding that the threshold shift of the animals from Oil/NE group was increased after hearing recovery suggests that noise exposure could accelerate the cochlear aging as a long-term effect after hearing recovery (Fernandez et al., 2015).

Moreeover, it was found that the mice that came from dams treated with oil and exposed later to high levels of flat noise (Oil/NE) were able to regain their hearing one week after noise exposure, indicated by a significant reduction of the hearing threshold shift on day 7 compared to day 0 for low-frequency pure tones (5 and 10 kHz) (Fig. 2E). In contrast, developmental exposure to the PCB mixture was found to prevent hearing recovery of the animals exposed later to high-level flat noise (PCB/NE) indicated by the nonsignificant reduction of the hearing threshold shift on day 7 compared to that on day 0 for all tested frequencies (Fig. 2E). Consistently, the threshold shift of the pups of the (Oil/NE) group on day 7 was significantly lower than that of the pups of the (PCB/NE) group for the low-frequency pure tones (5 and 10 kHz) (Fig. 2E -). Therefore, the above data suggest that developmental damage of the 6 mg/kg Fox River PCB mixture on hearing at flat noise and lower frequencies impairs the machinery of short-term hearing recovery after noise exposure later in adulthood. However, exposure to high levels of flat noise could accelerate cochlear aging as a long-term effect after hearing recovery.

### The effect of PCBs and/or noise exposure on the auditory midbrain

The hyperactivity of the IC was reported after acoustic trauma through many studies (Mulders and Robertson, 2009; Manzoor et al., 2012; Wei 2013; Gröschel et al., 2014), which could be associated with the development of tinnitus as a long-term effect after such acoustic insults, as reviewed (Berger and Coomber, 2015). Therefore, the IC is a good candidate to examine if developmental exposure to PCBs could exacerbate the long-term negative central effects of noise exposure later in adulthood. The IC comprises a major lemniscal division, the central nucleus (ICC), that receives primarily ascending auditory projections (Willard FH and DK, 1983; JF, 2001; DL, 2005), and two non-lemniscal divisions known as the dorsal (DCIC) and lateral (LCIC) cortices that receive massive descending auditory as well as multisensory inputs (Coleman and Clerici, 1987; Saldaña et al., 1996; Winer et al., 1998; Bajo and Moore, 2005; Schreiner and Winer, 2005; Loftus et al., 2008; Lesicko et al., 2016; Vaithiyalingam Chandra Sekaran et al., 2021). To examine the differential impact of PCBs on excitatory vs. inhibitory neurons in the IC, and because the IC is superficially accessible in mice (Fig. 3A), two-photon in vivo microscopy was used to monitor the evoked calcium signals of the DCIC neurons to pure tone stimuli. The calcium indicator jRGECO1a was delivered to the non-GABAergic and GABAergic neurons of the DCIC through a viral expression of the red calcium indicator driven by the *Syn* promoter after injecting the DCIC with adeno-associated virus (AAV) immediately after the final ABR measurement (post-noise-day 7). The expression of GFP in the GABAergic neurons of GAD67-GFP knock-in animals enabled us to distinctively monitor the activity of non-GABAergic (GFP-negative) and GABAergic (GFP-positive) neurons. Given that the chronic tinnitus-like behavior can take weeks or months after acoustic trauma to emerge (Longenecker and Galazyuk, 2011; Middleton et al., 2011; Turner et al., 2012), the acoustically evoked neuronal activity was imaged after 28 to 30 weeks of noise exposure to examine any possible long-term central effects. The recording of the evoked calcium signals to different (frequency/amplitude) combinations of pure tones was successfully achieved (Fig. 3B). Based on the best pure tone frequency of each responsive cell (Barnstedt et al., 2015), non-GABAergic and GABAergic cells of the DCIC of the animals from the Oil/NU group showed a similar tonotopic organization reported before (Barnstedt et al., 2015; Wong and Borst, 2019), which consisted of cells tuned to lower and higher frequencies along the rostromedial to the caudolateral axis, respectively. The positive correlation between the distance of each cell on the mediolateral axis and its best frequency is an indicator for the tonotopy shown by the Oil/NU group (Fig. 3C, 1^st^ row). Although PCB/NU and Oil/NE groups showed a significant positive correlation between the distance of the cells along the mediolateral axis and their best frequency indicating a good tonotopic organization over the mediolateral axis, both groups had different profiles of how the best frequencies of the cells are spatially distributed.While the Oil/NE group showed a similar tonotopic map to that of the Oil/NU group featuring some medially located cells tuned to mostly low-frequency pure tones(Fig. 3C: 3^rd^ rows), the PCB/NU group showed a high-low-high map featuring some medially located cells tuned to high-frequency pure tones (Fig. 3C: 2^nd^ row), which was previously reported (Wong and Borst, 2019). This difference in the tonotopic organization of the DCIC neurons across different exposures suggests that the tonotopy of the DCIC could be reshaped based on environmental input. The PCB/NE group showed a complete disruption of tonotopy (Fig. 3C: 4^th^ row) indicated by the lack of a correlation between the distance of the cells along the mediolateral axis and their best frequency. This disruption was characterized by a significant loss of the responsive non-GABAergic cells compared to all other groups (Fig. 3D). This non-responsive zone was mostly located in the lateral areas where the cells were tuned to the higher frequencies (Fig. 3C: 4^th^ row). Consistent with this finding, the distribution of the cells based on best frequency showed a significant low-frequency bias for GABAergic and non-GABAergic neurons in the PCB/NE group compared to all groups (Fig. 3E), which could indicate synaptic reorganization to compensate for the low-frequency hearing loss. A smaller effect was found in the Oil/NE group and PCB/NU groups (Fig. 3E), which could indicate a partial disruption of the DCIC tonotopy. Further, this loss of the neurons tuned to higher frequencies in the lateral areas of the DCIC of the PCB/NE group resulted in a complete reversal of tonotopy (higher to lower frequencies along the medial to the lateral axis) compared to all other groups (lower to higher frequencies along the medial to the lateral axis). However, it is not known if the change in the tonotopic organization of the DCIC is driven by central and/or peripheral components. The significantly higher threshold to flat noise of the animals from PCB/NU, Oil/NE, and PCB/NE groups at the time of the two-photon imaging compared to that of the control (Oil/NU) group (Fig. 2B) suggests that the changes in the DCIC induced by different exposures could be due to peripheral damage. Although PCB/NE and Oil/NE groups shared a similar threshold to the flat noise 4 months after noise exposure (Fig. 2B), PCB/NE group had a disrupted tonotopy compared to that shown by the Oil/NE group, which indicates that the DCIC changes induced by PCBs and noise exposure (PCB/NE) was accompanied by central changes.

**Figure 3.**
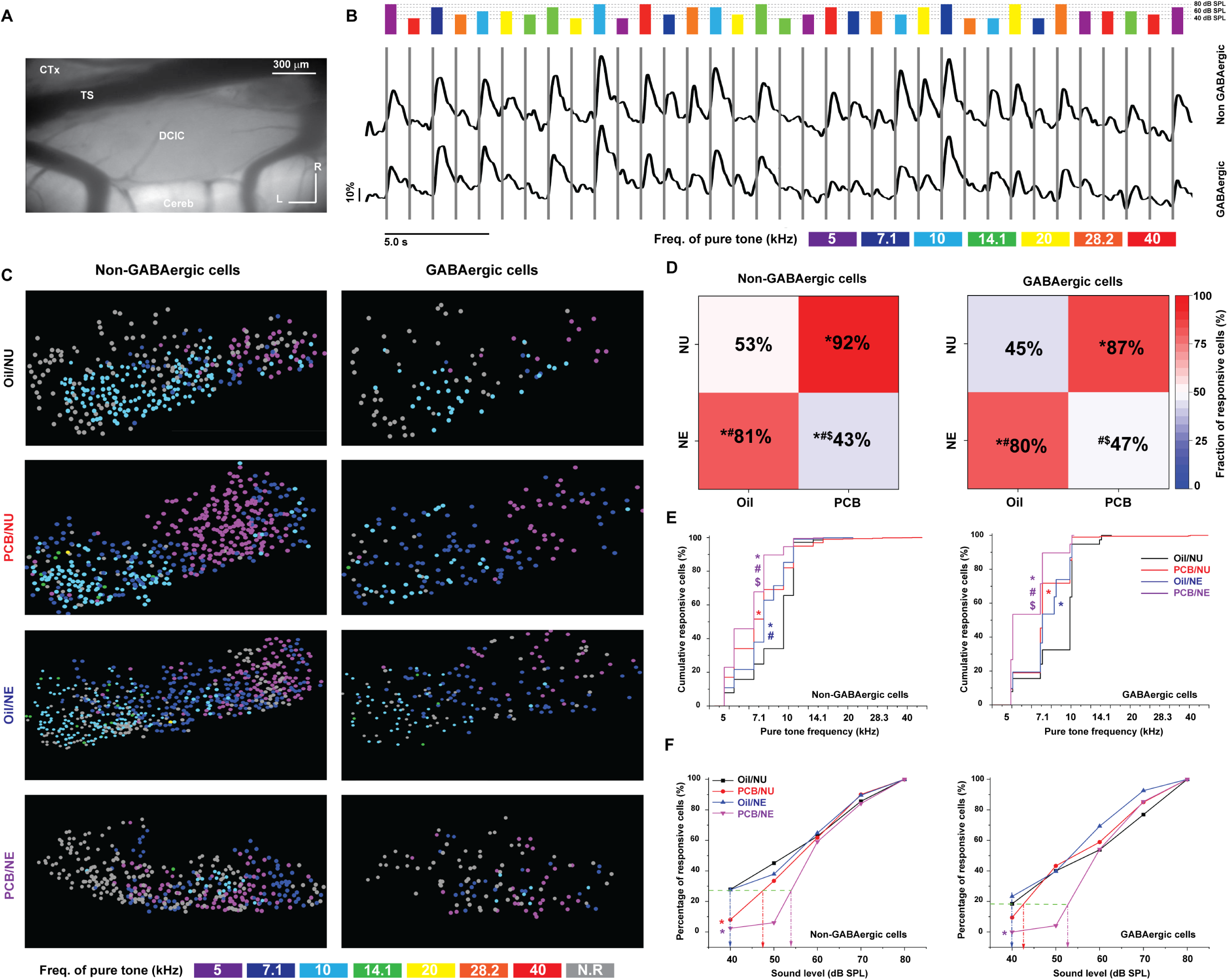
The effect of PCBs and/or noise exposure on the DCIC cells. ***A,*** A brightfield image of the surface of the DCIC. ***B,*** The time traces of the evoked calcium signals by different stimuli of pure tones obtained from non-GABAergic (top) and GABAergic (bottom) cells of the DCIC imaged from the surface; The length and the color of the gradient green bars indicate the intensity and the frequency of the stimulus, respectively. ***C,*** Pseudocolor images show the activity of the cells on the DCIC across different animals from (Oil/NU: 1^st^ row, PCB/NU: 2^nd^ row, Oil/NE: 3^rd^ row, and PCB/NE: 4^th^ row) in response to the pure tone based on their best frequency as graded green circles for the responsive cells and solid red circles for the non-responsive cells (Pearson Corr. 0.69 (p = 2.4×10^-37^), 0.56 (p = 2.6×10^-37^), 0.7 (p = 6.5×10^-69^), and −0.007 (p = 0.92) for non-GABAergic cells of Oil/NU, PCB/NU, Oil/NE, and PCB/NE, respectively & Pearson Corr. 0.63 (p = 1.9×10^-6^), 0.56 (p = 3.7×10^-13^), 0.62 (p = 5.6×10^-17^), and - 0.03 (p = 0.85) for GABAergic cells of Oil/NU, PCB/NU, Oil/NE, and PCB/NE, respectively). ***D,*** A heat map showing the fraction of the responsive non-GABAergic (left) (Chi-square test: χ^2^= 227 (*p <0.001), 111 (*p <0.001) and 9.9 (*p = 0.001) for Oil/NE vs.PCB/NU, Oil/NE, and PCB/NE, respectively & χ^2^= 36.3 (^#^p <0.001) and 311 (^#^p <0.001) for PCB/NU vs.Oil/NE and PCB/NE, respectively & χ^2^= 181 (^$^p <0.001) for Oil/NE vs.PCB/NE) and GABAergic (right) (Chi-square test: χ^2^= 79.8 (*p <0.001), 54.6 (*p <0.001) and 0.09 (p = 0.67) for Oil/NE vs.PCB/NU, Oil/NE, and PCB/NE, respectively & χ^2^= 4.2 (^#^p = 0.04) and 65.2 (^#^p <0.001) for PCB/NU vs.Oil/NE and PCB/NE, respectively & χ^2^= 42 (^$^p <0.001) for Oil/NE vs.PCB/NE) cells to sounds across all exposure groups. ***E,*** A cumulative distribution function for non-GABAergic (left) (Kruskal-Wallis ANOVA: χ^2^= 156, *p <0.001 for Oil/NE vs.PCB/NU, Oil/NE, and PCB/NE, respectively & ^#^p = 0.003 and ^#^p = 7.1×10^-7^ for PCB/NU vs.Oil/NE and PCB/NE, respectively & ^$^p <0.001 for Oil/NE vs.PCB/NE) and GABAergic (right) (Kruskal-Wallis ANOVA: χ^2^= 59.9, *p = 5.0×10^-6^, *p = 4.2×10^-4^, *p <0.001 for Oil/NE vs.PCB/NU, Oil/NE, and PCB/NE, respectively & p = 1.0 and ^#^p = 5.5×10^-5^ for PCB/NU vs.Oil/NE and PCB/NE, respectively & ^$^p = 7.0×10^-7^ for Oil/NE vs.PCB/NE) cells of the DCIC based on their best frequency. ***F,*** A line graph showing the fraction of evoked non-GABAergic (left) (Chi-square test: χ^2^= 53.3 (*p <0.001), χ^2^= 0.0043 (p = 0.99), and 43.6 (^#^p <0.001) for Oil/NU vs. PCB/NU, Oil/NE, and PCB/NE, respectively, at 40 dB SPL & χ^2^= 0.06 (p = 0.99), χ^2^= 0.34 (p = 0.95), and 0.52 (p = 0.91) for Oil/NU vs. PCB/NU, Oil/NE, and PCB/NE, respectively, at 60 dB SPL) and GABAergic (right) (Chi-square test: χ^2^= 3.7 (p = 0.29), χ^2^= 0.65 (p = 0.88), and χ^2^= 9.9 (*p = 0.02) for Oil/NU vs. PCB/NU, Oil/NE, and PCB/NE at 40 dB SPL & χ^2^= 0.42 (p = 0.92), χ^2^= 5.1 (p = 0.16), and 0.001 (p = 0.99) for Oil/NU vs. PCB/NU, Oil/NE, and PCB/NE, respectively, at 60 dB SPL) cells of the DCIC at each sound level within the responsive cell population across all exposure groups; The exposure groups are plotted as Oil/NU: black line, PCB/NU: red line, Oil/NE: blue line, and PCB/NE: purple line; *The significance against (Oil/NU) group; ^#^The significance against (PCB/NU) group; ^$^The significance against (Oil/NE) group; Cereb: Cerebellum; CTx: Cortex, DCIC: dorsal cortex of the inferior colliculus, TS: The transverse sinus. A rainbow color code was selected for different frequencies, N.R in grey: Non-responsive cells.

Compared to the Oil/NU group, both PCB/NU and Oil/NE groups had more responsive GABAergic and non-GABAergic neurons to pure tone stimuli (Fig. 3C: 2^nd^ and 3^rd^ rows & Fig. 3D), which could indicate an increased gain induced by homeostatic plasticity after the peripheral damage and consistent with the previous finding showing the increased gain of the corticocollicular axons reported after noise-induced hearing loss (Asokan et al., 2018). To test this possibility, the average cellular response across all tested frequencies and amplitudes was calculated based on the AUC of the evoked response. Consistent with our hypothesis, there was an increase in the evoked cellular response of non-GABAergic and GABAergic cells of the Oil/NE and PCB/NU groups compared to control (Oil/NU) (Non-GABAergic cells: Kruskal-Wallis ANOVA, χ^2^= 183.6, p = 1.49 x 10^-39^, post hoc Dunn’s Test: p = 1.5 x 10^-10^ or 5.1 x 10^-7^ for Oil/NE (Median = 270 .7) or PCB/NU (Median = 274.8) vs.Oil/NU (Median = 210.6), respectively & GABAergic cells: Kruskal-Wallis ANOVA, χ^2^= 66.7, p = 2.2 x 10^-14^, post hoc Dunn’s Test: p = 1.7 x 10^-7^ or 4.7 x 10^-6^ for Oil/NE (Median = 211 .5) or PCB/NU (Median = 228.9) vs.Oil/NU (Median = 105.2), respectively), indicating a higher gain of both cell types in Oil/NE and PCB/NU groups. In contrast, the response of non-GABAergic cells only of the PCB/NE group was reduced compared to the control (Non-GABAergic cells: Kruskal-Wallis ANOVA, χ^2^= 183.6, p = 1.49 x 10^-39^, post hoc Dunn’s Test: p = 1.95 x 10^-7^ for PCB/NE (Median = 142) vs.Oil/NU (Median = 210.6) & GABAergic cells: Kruskal-Wallis ANOVA, χ^2^= 66.7, p = 2.2 x 10^-14^, post hoc Dunn’s Test: p = 1.0 for PCB/NE (Median = 112.96) vs.Oil/NU (Median = 105.2)), suggesting that the combination between PCBs and noise exposure did not induce a synaptic compensatory mechanism to the peripheral damage. These results suggest that the cellular gain could be modulated based on the degree of the peripheral or central damage induced by different exposures in a frequency-dependent manner. Within the population of responsive neurons of the DCIC to sound in the unexposed control animals (Oil/NU), it was found that 27.5% and 18.3% of responsive non-GABAergic and GABAergic cells, respectively, could be activated by sound at 40 dB SPL. Therefore, we asked if a sound at 40 dB SPL could activate a similar percentage of cells in the DCIC of Oil/NE (hearing recovered) vs.PCB/NE (hearing unrecovered). Whereas a similar percentage of non-GABAergic and GABAergic cells could be activated by a 40 dB SPL sound level in the Oil/NE compared to the Oil/NU group (Fig. 3F: 27.6% and 23.3% of responsive non-GABAergic and GABAergic cells, respectively), a significantly lower percentage of cells were activated by a 40 dB sound level in the PCB/NE group (Fig. 3F: 2.4% and 0.0% of responsive non-GABAergic and GABAergic cells, respectively). By interpolation, the PCB/NE group was found to require a higher sound level (53.8 and 52.6 dB SPL) to activate a similar fraction of the responsive non-GABAergic and GABAergic cells, respectively (Fig. 3F) as the Oil/NU group, which was consistent with the persistent toxic effect of PCBs that prevented the hearing recovery. Consistent with the hearing damage induced by PCB exposure, the DCIC cells of PCB/NU showed a significantly lower fraction of activated non-GABAergic cells by a 40 dB SPL sound level compared to those of the Oil/NU group (Fig. 3F: 8.0% and 9.4% of responsive non-GABAergic and GABAergic cells, respectively) and then required a moderately higher sound level (47.6 and 42.5 dB SPL) to activate a similar percentage of non-GABAergic and GABAergic cells, respectively, compared to the Oil/NU group (Fig. 3F). In addition, all groups showed a similar fraction of activated cells at a 60 dB SPL sound level, suggesting that most differences were seen at relatively low sound pressure levels (Fig. 3F). The above data show that the combination of developmental exposure to PCBs and noise exposure in adulthood had a significant long-term negative impact on the DCIC neurons, which is characterized by a disruption of the tonotopic organization and an increase of the threshold of DCIC neurons.

### The effect of PCBs and/or noise exposure on the inhibition and excitation balance of the DCIC

For each amplitude/frequency combination, the pooled AUC values of the calcium signals obtained from either non-GABAergic or GABAergic cells were calculated to examine excitatory or inhibitory activities, respectively, of the DCIC across different exposures. After normalizing the AUC values against the highest value across all amplitude/frequency combinations per each animal, a heat map was made by averaging the AUC values across all animals in each group. The visual assessment of these maps indicated that the frequency response areas (FRAs) in the exposure groups (PCB/NU, Oil/NE, and PCB/NE) were smaller compared to the control group (Oil/NU). To quantify this difference, the sound level evoking 20% of the highest response at each frequency was assigned as a sound threshold at that frequency for each animal within each group. Compared to the control group, all animals exposed to either PCBs, noise, or their combination required a significantly higher sound pressure level to get a 20% of response at 10 and 14.1 kHz for non-GABAergic cells and GABAergic cells (Fig. 4B and C, Table 4). The PCB-exposed animals either those exposed to PCBs alone or PCBs combined with noise required a significantly higher sound level to achieve 20% activity of their GABAergic cells at 28.3 kHz (Fig. 4C, Table 4). As indicated by the visual assessment of the heat maps, the FRA of inhibitory or GABAergic cells of the PCB/NE group was smaller than that of the control group (Oil/NU), which indicated a loss of inhibition induced by PCB exposure. Given that this loss of inhibition could be associated with a loss of excitation of a similar degree, the inhibition/excitation (INH/EXC) ratio was computed to examine the occurrence of inhibitory downregulation. The (INH/EXC) ratio was calculated by dividing the pooled AUC values of GABAergic by those of non-GABAergic cells at either each sound amplitude across different frequencies or at each sound frequency across different amplitudes. Although the (INH/EXC) ratio was not significantly different between all groups at all sound amplitudes (Fig. 4D), the (INH/EXC) ratio of all exposure groups was significantly lower than that of the control group (Oil/NU) at 5 kHz (Fig. 4E). In addition, the combination of PCBs and noise exposure (PCB/NE) resulted in a persistent low (INH/EXC) ratio at 7.1 kHz compared to the control group (Oil/NU). Also, the noise-exposed groups had a lower (INH/EXC) ratio than that of (PCB/NU) at 7.1 kHz (Fig. 4E). These data suggest that PCBs, noise, or their combination could manipulate the activity of the DCIC in a frequency-dependent manner. For instance, at higher frequencies (10-14 kHz), the PCBs and/or noise exposure increase the threshold of both excitatory and inhibitory activities in a balanced manner indicated by no change in the (INH/EXC) ratio. In contrast, the PCBs and/or noise exposure disrupted the balance between inhibition and excitation by downregulating the inhibition at 5 kHz with no change in the activity threshold. However, only the combination of PCBs and noise exposure extended the inhibitory downregulation at 7.1 kHz.

**Figure 4.**
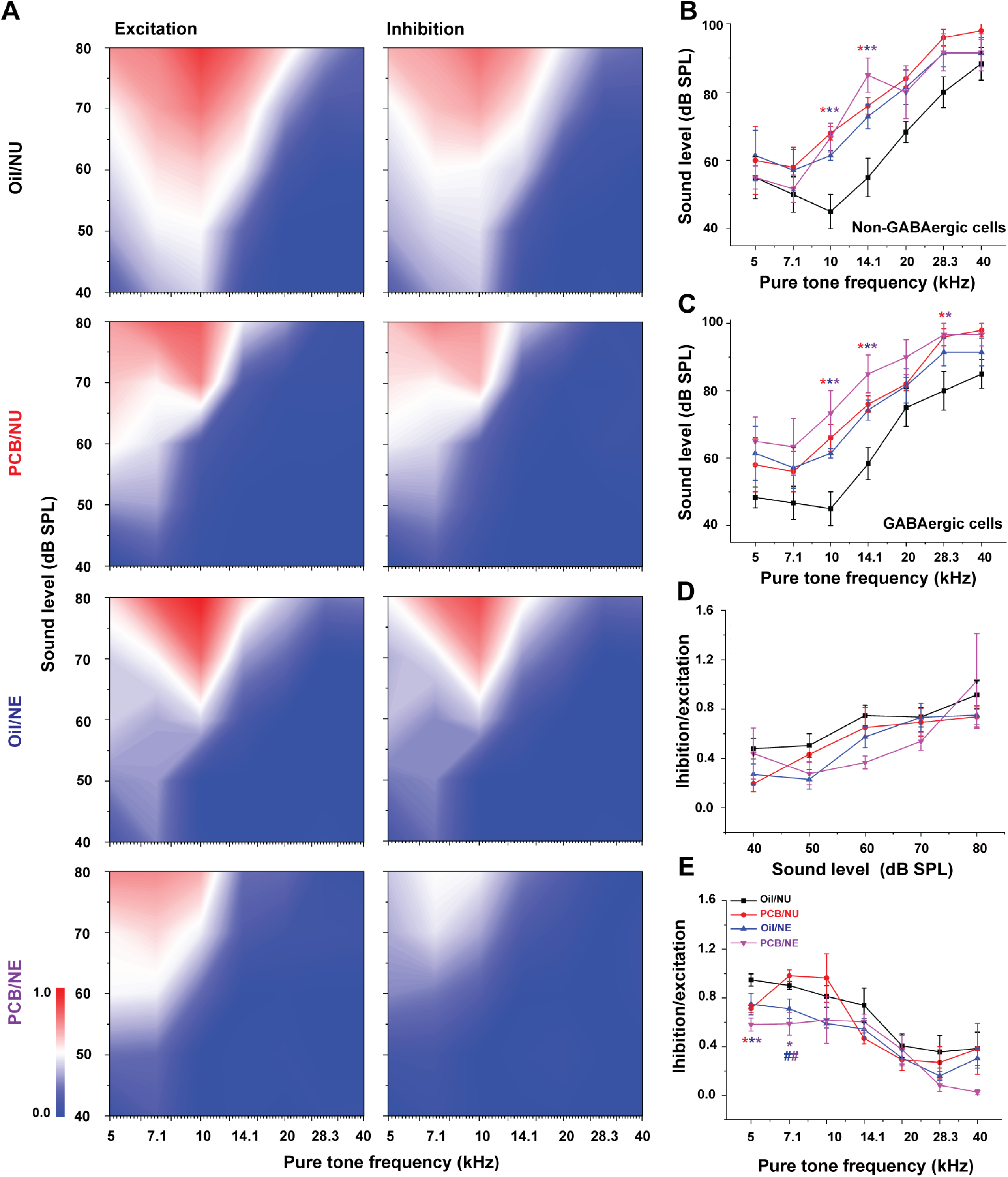
The effect of PCBs and/or noise exposure on the inhibition and excitation balance of the DCIC. ***A,*** Heat maps showing the excitatory (left) and inhibitory (right) average response across the animals of each exposure group (Oil/NU: 1^st^ row, PCB/NU: 2^nd^ row, Oil/NE: 3^rd^ row, and PCB/NE: 4^th^ row). ***B-C,*** Line graphs showing the sound level required to evoke 20% of the sound response in non-GABAergic (One way ANOVA: At 10 kHz, f _(3, 20)_ = 9.0, p = 5.7×10^-4^, Fisher post hoc test: *p = 0.0002, 0.002, and 0.0002 for Oil/NU vs.PCB/NU, Oil/NE, and PCB/NE, respectively, p = 0.2 for PCB/NU vs. Oil/NE, 0.79 for PCB/NU vs. PCB/NE, and 0.28 for Oil/NE vs. PCB/NE & at 14.1 kHz, f _(3, 20)_ = 8.0, p = 0.001, Fisher post hoc test: *p = 0.004, 0.008, and 0.0001for Oil/NU vs.PCB/NU, Oil/NE, and PCB/NE, respectively, p = 0.62 for PCB/NU vs. Oil/NE, 0.18 for PCB/NU vs. PCB/NE, and 0.06 for Oil/NE vs. PCB/NE & f _(3, 20)_ = 0.24, p = 0.86 at 5kHz, f _(3, 20)_ = 0.53, p = 0.67 at 7.1 kHz, f _(3, 20)_ = 1.7, p = 0.19 at 20 kHz, f _(3, 20)_ = 2.3, p = 0.1 at 28.3 kHz, and f _(3, 20)_ = 0.75, p = 0.35 at 40 kHz) or GABAergic (One way ANOVA: At 10 kHz, f _(3, 20)_ = 6.9, p = 0.002, Fisher post hoc test: *p = 0.006, 0.02, and 0.0003 for Oil/NU vs.PCB/NU, Oil/NE, and PCB/NE, respectively, p = 0.49 for PCB/NU vs. Oil/NE, 0.29 for PCB/NU vs. PCB/NE, and 0.07 for Oil/NE vs. PCB/NE & at 14.1 kHz, f _(3, 20)_ = 6.9, p = 0.002, Fisher post hoc test: *p = 0.01, 0.01, and 0.0002 for Oil/NU vs.PCB/NU, Oil/NE, and PCB/NE, respectively, p = 0.78 for PCB/NU vs. Oil/NE, 0.16 for PCB/NU vs. PCB/NE, and 0.08 for Oil/NE vs. PCB/NE & at 28.3 kHz, f _(3, 20)_ = 3.3, p = 0.04, Fisher post hoc test: *p = 0.02, p = 0.06 and *p = 0.01 for Oil/NU vs.PCB/NU, Oil/NE, and PCB/NE, respectively, p = 0.46 for PCB/NU vs. Oil/NE, 0.92 for PCB/NU vs. PCB/NE, and 0.37 for Oil/NE vs. PCB/NE & f _(3, 20)_ = 1.1, p = 0.38 at 5kHz, f _(3, 20)_ = 1.1, p = 0.36 at 7.1 kHz, f _(3, 20)_ = 1.6, p = 0.23 at 20 kHz, f _(3, 20)_ = 0.75, p = 0.35 at 40 kHz) cells, respectively, across different pure tone frequencies for each group. ***D-E,*** Line graphs showing the inhibition/excitation ratio across different amplitudes or frequencies (One way ANOVA: At 5 kHz, f _(3, 18)_ = 4.9, p = 0.01, Fisher post hoc test: *p = 0.037, 0.039, and 0.001 for Oil/NU vs.PCB/NU, Oil/NE, and PCB/NE, respectively, p = 0.73 for PCB/NU vs. Oil/NE, 0.23 for PCB/NU vs. PCB/NE, and 0.09 for Oil/NE vs. PCB/NE & at 7.1 kHz, f _(3, 16)_ = 5.4, p = 0.009, Fisher post hoc test: *p = 0.01 for Oil/NU vs.PCB/NU, ^#^p = 0.02 and 0.003 for PCB/NU vs.Oil/NE and PCB/NE, respectively, p = 0.47 for Oil/NU vs. PCB/NU, 0.06 for Oil/NU vs. Oil/NE, and 0.24 for Oil/NE vs. PCB/NE & f _(3, 19)_ = 1.8, p = 0.18 at 10kHz, f _(3, 19)_ = 1.4, p = 0.27 at 14.1 kHz, f _(3, 18)_ = 0.36, p = 0.78 at 20 kHz, f _(3, 17)_ = 1.4, p = 0.29 at 28.3 kHz, and f _(3, 17)_ = 1.3, p = 0.30 at 40 kHz), respectively, of pure tone stimulus for each exposure group; The exposure groups are plotted as Oil/NU: black line, PCB/NU: red line, Oil/NE: blue line, and PCB/NE: purple line; *The significance against (Oil/NU) group; ^#^The significance against (PCB/NU) group.

### The association between the downregulation of inhibition and oxidative stress

Given that both PCB and noise exposure are associated with high levels of oxidative stress (Spreng, 2000; Samson et al., 2008; Bavithra et al., 2012; Lee et al., 2012; Selvakumar et al., 2012; Daiber et al., 2020; Mao and Chen, 2021; McCann et al., 2021), it was important to examine the capacity of the animals developmentally exposed to PCBs and/or noise in adulthood to mitigate oxidative stress. Therefore, the brains of the animals across different exposures were isolated at different time points (three months of age after the last ABR measurement which was post-noise-day 7 and seven months of age after the two-photon imaging). The brains were frozen and sectioned at the level of the IC. Then, multiple 1 mm biopsies were taken from the IC structure using a micro-puncture (Fig. 5A). The tissue specimens were processed for Western Blot to examine the levels of the metabolic enzyme sodium dismutase-2 (SOD2), which is one of the important enzymes that function to scavenge reactive oxygen species (ROS) (Fig. 5B). Although the levels of SOD2 showed no significant differences between the exposure groups, low SOD2 levels were found in older animals of higher body weight that were exposed to PCBs and/or noise indicated by a significant negative correlation between the SOD2 levels and the age or the body weight of the animals. This effect was stronger in the PCB/NE group (Figs. 5C and D). This finding was consistent with many reports showing that aging or higher body weight could induce higher levels of oxidative stress that could be associated with lower levels of SOD2 (Pansarasa et al., 1999; Tatone et al., 2006; Zeng et al., 2014; Balasubramanian et al., 2020). Moreover, consistent with the strong correlation between noise exposure and high levels of oxidative stress, low SOD2 levels were associated with higher hearing threshold values in the Oil/NE group only (Fig. 5E). GABAergic cells are known for their vulnerability to oxidative stress (as reviewed (Ibrahim and Llano, 2019)), so we asked if the downregulation of inhibition induced by PCBs and noise exposure is associated with high levels of oxidative stress. Therefore, the tissue specimens were processed for Western Blot to examine the levels of glutamate acid decarboxylase-67 (GAD67), which is the main cytosolic enzyme to produce GABA as an indicator for the level of inhibition, and SOD2 (Fig. 5B). Initially, the GAD67 levels showed no significant difference between the exposure groups, which was consistent with the previous report (Bandara et al., 2016). However, only the PCB/NE group showed a significant positive correlation between GAD67 and SOD2 (Fig. 5F) suggesting that GABAergic inhibition was retained in animals with a higher capacity to clear the ROS, which may suggest that at least part of the downregulation of inhibition seen after PCB and noise exposure may be related to an increase in oxidative stress in the IC.

**Figure 5.**
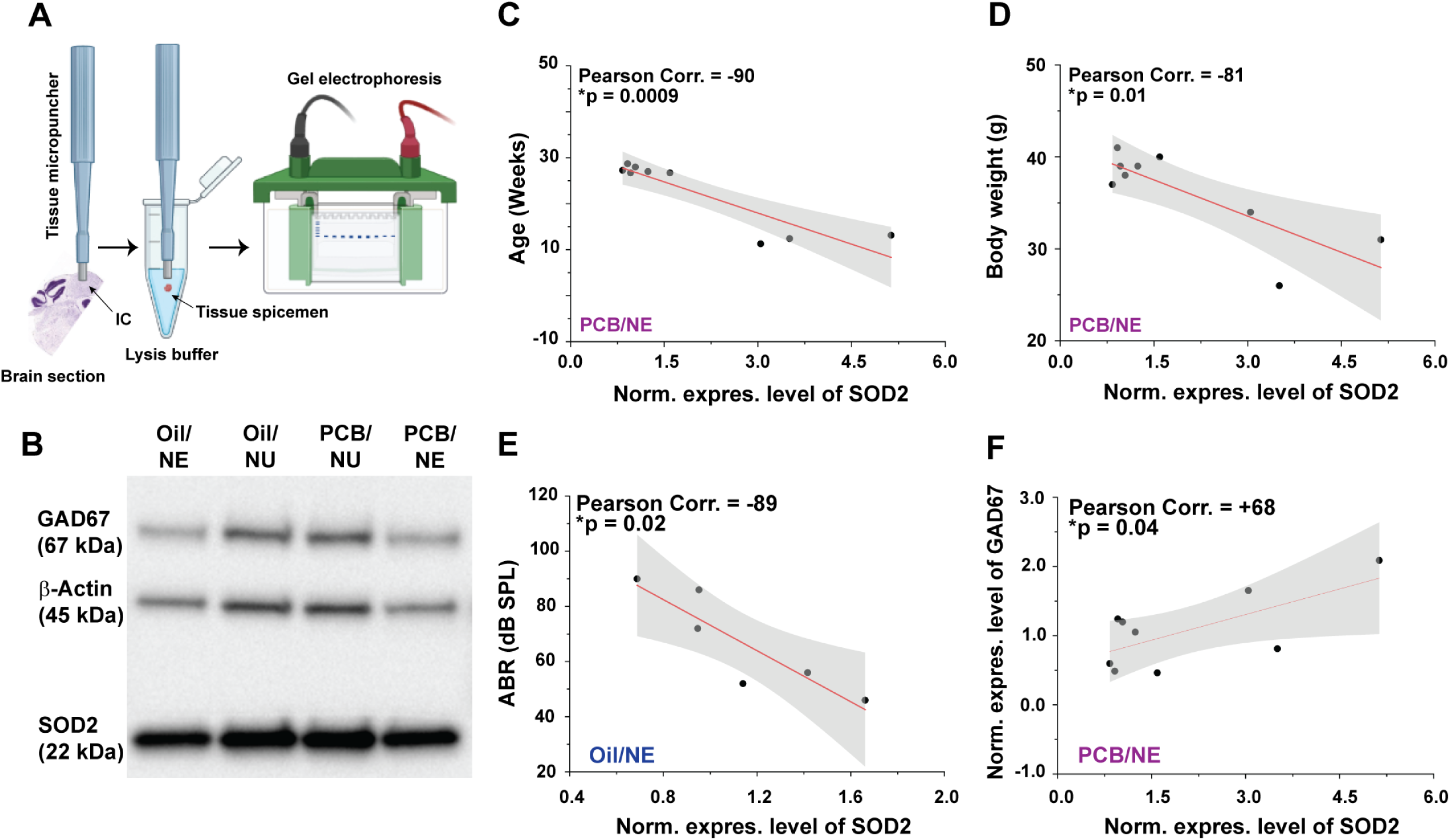
The association between the downregulation of inhibition and oxidative stress. ***A,*** A cartoon image showing the process of tissue collection and gel running for Western Blot (Some features were taken from https://biorender.com). ***B,*** An image showing the Western Blots of GAD67 (1^st^ row), actin (2^nd^ row), and SOD2 (3^rd^ row). ***C-F,*** Line graphs showing the correlation between the normalized expression levels of the SOD2 enzyme and either age, body weight, or the normalized expression level of the GAD67 enzyme for the (PCB/NE) group as well as ABR for the (Oil/NE) group. *: p-values for each correlation.

## Discussion

The current study shows that developmental exposure to PCBs resulted in a sex-dependent low-frequency hearing loss in mice, that pre- and perinatal PCB exposure prevented the normal recovery of hearing after adult noise exposure, that PCB and noise exposure leads to downregulation of synaptic inhibition and disruption of tonotopy of the IC, and that this downregulation of inhibition may be related to increases in oxidative stress. Thus, this work provides a new mechanistic understanding of how an environmentally common combination of toxins leads to central changes in the auditory system and how developmental toxin exposure can have long-lasting neural consequences. Below we address the technical considerations and implications of this work.

### Technical considerations and comparison with previous work

The female mice exposed to PCBs and the breeding GAD67-GFP knock-in male mice share the same SW background. The SW is a well-known outbred strain that has generally good hearing into young adulthood (Zheng et al., 1999). Consistent with that, the control animals showed no change in their hearing threshold throughout the study (7 months of age) (Fig. 2B) indicating limited aging or genetic-related negative effects on hearing during the study. A relatively short duration/high intensity noise exposure paradigm was used in this study. This paradigm was used because previous work had shown that rats developmentally exposed to PCBs developed audiogenic seizures (AGS) after they were exposed to higher sound pressure levels (Poon et al., 2015; Bandara et al., 2016). Therefore, in our paradigm, mice were anesthetized during noise exposure, which limits the activity of the middle ear reflex (MER), which is known for its protective role against acoustic damage (Brask, 1979; Deiters et al., 2019). Thus, the particular noise exposure protocol here was more likely to cause hearing loss than in awake animals, as reflected in the increased hearing thresholds seen in this study.

Two-photon imaging of the IC was conducted under ketamine anesthesia. As a well-known anesthetic, ketamine has been used in some studies showing the large-scale tonotopic organization in the mouse auditory cortex (Stiebler et al., 1997; Linden et al., 2003; Guo et al., 2012; Yudintsev et al., 2021), which showed a similar topographical functional organization under ketamine and awake state (Guo et al., 2012). However, ketamine may broaden the bandwidth of the FRA and lengthen the response duration (Guo et al., 2012), which could impact the tonal selectivity and temporal properties of the responses. Also, ketamine was reported to increase the spontaneous activity of the IC cells in guinea pigs (Astl et al., 1996; Syka et al., 2000) and the auditory cortex of the rats, which resulted in trial-to-trial variability (Kisley and Gerstein, 1999). To correct for such variability between runs, the average response of all runs across the tested amplitudes at each frequency was taken. The same procedures were followed across all exposure groups.

Our results showed that the difference between the hearing thresholds and impact of PCB exposure in phase 2 (Fig. 2C) was larger than that found in phase 1 (Fig. 1C) across the tested frequencies. The cause of these cohort-effects are not known, though similar cohort effects have been seen in previous developmental PCB exposure work (Lee et al., 2021). In the current study, phase 1 and phase 2 were performed one year apart with separate outbred SW breeders purchased for each phase, suggesting that genetic variability may contribute to these differences (Cui et al., 1993). The differences seen in the two cohorts reinforce the need to use matched cohorts for PCB-based studies, particularly in outbred strains. However, despite these differences, the trends seen in both cohorts are highly consistent with previous studies establishing PCB effects on hearing (Goldey et al., 1995; Crofton and Rice, 1999; Crofton et al., 2000; Powers et al., 2006; Powers et al., 2009).

Our results showed that the low INH/EXC ratio found in PCB/NE observed via two-photon imaging was associated with no change in the expression level of GAD67 across all groups, while the previous study done on rats (Bandara et al., 2016) showed a decrease in IC GAD expression after PCB exposure. Such contradictory results could be explained by the difference in the experimental sampling and scaling. While the two-photon imaging was done on the DCIC, the whole IC tissue was dissected for protein quantification due to the technical difficulty to obtain a pure representative sample of the DCIC. Therefore, the results obtained by each technique could implicate sub-region-specific changes in IC after the acoustic trauma. Consistent with that notion, and indicated by the click-evoked potentials measured from the external cortex of the IC and its central nucleus after acoustic trauma, it was found that the responses of the two IC sub-regions are different (Szczepaniak and Møller, 1996). Such differences could originate from the difference in input and output between the lemniscal (Willard FH and DK, 1983; JF, 2001; DL, 2005) and non-lemniscal (Coleman and Clerici, 1987; Saldaña et al., 1996; Winer et al., 1998; Bajo and Moore, 2005; Schreiner and Winer, 2005; Loftus et al., 2008) regions of the IC.

### Implications

#### PCBs, hearing, and sex differences

Our data are consistent with multiple previous studies demonstrating that developmental PCB exposure causes hearing loss (Goldey et al., 1995; Herr et al., 1996; Crofton and Rice, 1999; Crofton et al., 2000; Lasky et al., 2002; Powers et al., 2006; Kenet et al., 2007; Powers et al., 2009). Several hypotheses exist to explain the impact of developmental PCB exposure on hearing. First, the toxic effect of PCBs on the thyroid gland leads to developmental alteration to the cochlea (Uziel, 1986; Goldey et al., 1995; Goldey and Crofton, 1998; Knipper et al., 2000; Song et al., 2008). Other studies showed the direct toxicity of PCBs on the development of the cochlea by allowing elevations of intracellular calcium concentration in the outer hair cells via ryanodine receptors (Wong et al., 1997; Bobbin, 2002; Sziklai, 2004). Our data also showed that the PCBs and/or noise exposure was associated with long-term permanent damage to hearing, which seemed to be uniquely driven by those exposures without aging-related effects at least four months after acoustic trauma or seven months of age, which is consistent with other reports showing hearing impairment at relevant time points (Powers et al., 2009; Liu et al., 2018). Also, the data showed that noise exposure can accelerate aging associated hearing loss after short-term hearing recovery, which was consistent with previous work (Fernandez et al., 2015).

Regarding the sex-dependent toxic effect of PCB exposure, previous studies reported that developmental exposure to PCBs showed mixed sex-dependent effects on behavior, locomotion, and vision in rats (Kremer et al., 1999; Geller et al., 2001; Vega-López et al., 2007; Cauli et al., 2013; Bell et al., 2016). Here, low-frequency hearing loss induced by developmental exposure to PCBs was only shown in male offspring. This finding could implicate the possible protective role of estrogen against hearing loss. For example, postmenopausal women were found to have a higher ABR threshold compared to younger women or men (Wharton and Church, 1990). Similar increases in thresholds were found in ovariectomized rats compared to the controls (Coleman et al., 1994). Estrogen was also found to have a protective effect against cochlear injury, gentamicin ototoxicity, noise, or age-related hearing loss (Nakamagoe et al., 2010; Tabuchi et al., 2011; Caras, 2013; Milon et al., 2018).

#### PCBs, noise, and the central auditory system

In addition to its negative impact on the peripheral auditory system, many studies examined the central effects of developmental exposure to PCBs or NIHL. The reduction of GABAergic inhibition (Poon et al., 2015; Bandara et al., 2016; Lee et al., 2021) and tonotopic reorganization (Kenet et al., 2007; Izquierdo et al., 2008; Pienkowski and Eggermont, 2009) are the most central common effects of PCBs and noise exposure. Consistent with previous reports, the DCIC of control animals showed a tonotopic map of non-GABAergic and GABAergic cells along the mediolateral axis (Barnstedt et al., 2015; Wong and Borst, 2019). Although similar distributions of responsive cells were seen after PCB exposure or noise exposure, there were more responsive cells to sound than control (Figs. 3C and D), which could be due to compensatory plasticity (Wang et al., 1996; Mulders and Robertson, 2009; Niu et al., 2013; Auerbach et al., 2014; Asokan et al., 2018). After PCB or noise exposure, the tuning frequencies of the cells were biased toward low frequencies (Fig. 3E), which could represent a compensatory mechanism in response to low-frequency hearing loss. In addition, the disruption of the tonotopic organization of the DCIC following the combination of PCBs and noise exposure involved both non-GABAergic and GABAergic cell populations. This tonotopic disruption was characterized by low responsiveness to sound and a possibility of reversed tonotopy of cells. The lower fraction of the responsive cells to sound in the PCB/NE group was only significant within the non-GABAergic neuronal population compared to the control group, which could indicate a selective synaptic mechanism targeting those cells. Our data are consistent with previous work showing that the PCB exposure disrupts the tonotopy of the primary AC, which was characterized by a non-responsive zone to sound or a reversed tonotopy (Kenet et al., 2007). Given that different exposure groups had higher hearing thresholds compared to the control at the time of two-photon imaging (4 months after noise exposure), the changes in the DCIC could be related to peripheral hearing loss, which is well-established to cause changes at the level of the IC (Ma et al., 2006; Izquierdo et al., 2008; Mulders and Robertson, 2011; Manzoor et al., 2012; Gröschel et al., 2014; Heeringa and van Dijk, 2014; Vogler et al., 2014). However, noise-exposed animals with or without PCBs exposure had similar hearing thresholds at the time of imaging, while the DCIC of the PCB/NE group showed a disrupted tonotopic organization. These data suggest that collicular changes induced by the combination between PCBs and noise exposure lead to central reorganization independent of peripheral hearing loss. For instance, the presence of tonotopic disruption at the level of the midbrain suggests that disruption in the AC may be inherited from lower centers. Alternatively, given the massive feedback to the DCIC from the AC (Asilador and Llano, 2020; Vaithiyalingam Chandra Sekaran et al., 2021; Yudintsev et al., 2021), the tonotopic disruption observed in the current study could be related to disrupted top-down corticocollicular inputs. Future work will be needed to sort out these possibilities.

#### PCBs, noise, GABA, and oxidative stress

PCB exposure in rats was associated with audiogenic seizures, which were associated with the downregulation of the inhibition indicated by the lower levels of the GAD56 enzyme (Bandara et al., 2016). Also, the NIHL causes neural hyperactivity in the central auditory system as reviewed (Zhao et al., 2016). Therefore, it was important to examine the balance between excitation and inhibition following PCBs exposure and/or noise exposure. The current study demonstrated that the PCB/NE group showed the smallest FRA for inhibitory neurons. It was also found that all PCB-exposed animals and/or high-level noise required a higher sound level to reach 20% of its response in both GABAergic and non-GABAergic cell populations. Moreover, a lower INH/EXC ratio was shown by all exposure groups at 5 kHz and by PCB/NE at 7.1 kHz, which suggests that the disruption of the balance between the excitation and inhibition by those exposures could be frequency-dependent. These results were supported by previous findings of PCB-induced disruption of the dynamics between the excitatory and inhibitory neurotransmission in the AC (Kenet et al., 2007; Lee et al., 2021). At the level of the IC, PCBs exposure was found to decrease the inhibition by reducing the level of GAD65 expression (Bandara et al., 2016). In addition, noise-induced hearing loss was found to disrupt the balance between inhibition and excitation in the IC and AC in mice (Scholl and Wehr, 2008; Roberts et al., 2010; Wang et al., 2011; Sturm et al., 2017).

Many GABAergic cells lack adequate defense mechanisms against oxidative stress (Ibrahim and Llano, 2019), which makes them more vulnerable to oxidative stress compared to other types of neurons. Given that PCBs and noise exposure are linked with an increase in oxidative and metabolic stress (McCann et al., 2021) (Pessah et al., 2019; Daiber et al., 2020; Liu et al., 2020), the downregulation of inhibition could be associated with a higher level of oxidative stress. The IC is among the most metabolically active regions in the brain (Landau et al., 1955; Sokoloff, 1981; Gross et al., 1987; Zeller et al., 1997; Bordia and Zahr, 2020). Therefore, we examined the association between the downregulation of inhibition and the oxidative stress indicated by the expression levels of both GAD67 and SOD2, respectively. As expected, the exposure to PCBs and/or noise promoted a positive correlation between oxidative stress, as indicated by the high levels of SOD2, and age and weight gain, which was consistent with previous reports (Pansarasa et al., 1999; Tatone et al., 2006; Zeng et al., 2014; Balasubramanian et al., 2020). Despite the absence of change in the level of both GAD67 and SOD2 levels across groups, a positive correlation between GAD67 and SOD2 expression was found in the PCB/NE group only, suggesting that GABAergic inhibition was retained in animals with a higher capacity to clear ROS. Indeed, future studies will be required to investigate the expression levels of different components involved in GABA production and/or function such as GAD65, which was reported to be reduced by PCB exposure in rats (Bandara et al., 2016), and GABA receptors. In addition, regional heterogeneity to oxidative stress, possibly related to perineuronal net expression, may lead to differential sensitivity to PCBs and noise based on the IC subregion (Schofield and Beebe, 2019).

#### Interactions between PCBs and noise exposure

Most of the effects of combined exposure to PCBs and noise were not entirely predictable by the effects of exposure to either PCBs or noise individually. For example, PCBs and noise exposure appear to combine nonlinearly to produce peripheral hearing loss. This interaction is evidenced by the finding that developmental exposure to PCBs did not exacerbate acute NIHL, but it prevented hearing recovery after one week of the acoustic trauma (Figure 2). Centrally at the level of the DCIC, while the individual exposures to PCBs or noise increased the number of responsive cells and their gain, the combined exposure to PCBs and noise reversed those effects (Fig. 3F). However, some linear interactions were also shown. These nonlinear interactions emphasize the need to conduct studies that examine the impact of both individual and combined toxin exposure on the developing auditory system.

In conclusion, developmental exposure to PCBs was found to impair the hearing of the male offspring of PCB-treated mouse dams and to prevent hearing recovery after acoustic trauma. PCBs exposure and noise exposure resulted in the disruption of the tonotopic map of the DCIC that was characterized by a wide non-responsive zone to sound, reversed, and low-frequency-biased tonotopy. Such effects were associated with the disruption of the excitation and inhibition balance via the downregulation of inhibition at lower frequencies and associated with diminished capacity to reduce oxidative stress. Given the level of PCBs still present in the environment, and the high (and growing) incidence of low-level noise exposure in our society (Beyer and Biziuk, 2009; Flamme et al., 2012; Lie et al., 2016; Weber et al., 2018), the current data in combination with previous work suggest that the auditory consequences of these exposures are likely widespread, synergistic, sex-dependent and involve disruption to broad areas of the central auditory system. Future work will be needed to understand the mechanisms by which these changes occur so that approaches may be designed to mitigate the damage caused by these common environmental toxicants.

## Acknowledgments

The authors deeply thank the National Institute of Environmental Health Sciences (NIEHS) for funding this work under the NIH Postdoctoral trainee in Endocrine, Developmental & Reproductive Toxicology (T32 ES007326), the National Instititute on Deafness and other Communication Disorders (R01 DC016599) and the NIH Office of Research Infrastructure Programs (S10 OD023569).

